# Personalized Cancer Therapy Prioritization Based on Driver Alteration Co-occurrence Patterns

**DOI:** 10.1101/772673

**Authors:** Lidia Mateo, Miquel Duran-Frigola, Albert Gris-Oliver, Marta Palafox, Maurizio Scaltriti, Pedram Razavi, Sarat Chandarlapaty, Joaquin Arribas, Meritxell Bellet, Violeta Serra, Patrick Aloy

## Abstract

Identification of actionable genomic vulnerabilities is the cornerstone of precision oncology. Based on a large-scale drug screening in patient derived-xenografts, we uncover connections between driver gene alterations, derive Driver Co-Occurrence (DCO) networks, and relate these to drug sensitivity. Our collection of 53 drug response predictors attained an average balanced accuracy of 58% in a cross-validation setting, which rose to a 66% for the subset of high-confidence predictions. Morevover, we experimentally validated 12 out of 14 *de novo* predictions in mice. Finally, we adapted our strategy to obtain drug-response models from patients’ progression free survival data. By revealing unexpected links between oncogenic alterations, our strategy can increase the clinical impact of genomic profiling.

## Background

In light of the complexity and molecular heterogeneity of tumors, clinical and histopathological evaluation of cancer patients is nowadays complemented with genomic information. Genome-guided therapy has been shown to improve patient outcome [1, 2] and clinical trial success rate [3] and, despite some controversy [4], prospective molecular profiling of personal cancer genomes has enabled the identification of an increasing number of actionable vulnerabilities [5].

Cancer genome sequencing initiatives have found that any given tumor contains from tens to thousands of mutations. However, only a few of them confer a growth advantage to cancer cells, driving thus the tumorigenic process. The most comprehensive study of ‘driver’ genes published to date has analyzed over 9,000 tumor samples, across 33 tissues of origin, and has systematically identified driver mutations in 258 genes [6]. Approximately half (142) of those driver genes were associated with a single tumor type, whereas 87 genes seem to provide a growth advantage in several tumor types. The number of drivers detected per tumor type varies widely, ranging from 2 in kidney chromophobe cancer to 55 in uterine cancer. Despite the large number of drivers identified per tumor type, every patient has a unique combination of mutations and copy number variants: ninety percent of patients show at least one putative driver alteration, but each sample only contains a median of three putatively altered drivers [7].

On top of identifying key alterations in tumor development, it is fundamental to pinpoint those that can shed light on the most appropriate therapy to treat each tumor (i.e. biomarkers). Often, patients with similar clinicopathological characteristics might be molecularly different [6], this inter-patient heterogeneity is one of the reasons why only a subset of them will actually respond to a given targeted treatment. Computational studies suggest that up to 90% of patients may benefit from molecularly-guided therapy when biomarkers of uncertain clinical significance, as well as off-label and experimental drugs, are used to guide treatment selection [7, 8]. Although randomized controlled trials are still considered the gold standard in the clinics, they cannot address all possible patient clinicopathologic and molecular subtypes [9]. Precision medicine has prompted the reconsideration of clinical drug development pipelines, with the implementation of more sophisticated clinical trial designs, such as umbrella, basket, and platform trials to account for inter-patient heterogeneity [10]. In particular, the implementation of adaptive enrichment strategies allows for continual learning and modification of the eligibility criteria as data accumulate, with the objective of recruiting those patients that are most likely to benefit from treatment [9–12]. However, despite the implementation of these novel experimental designs, currently only alterations in 28 genes have accumulated enough clinical evidence to be approved as biomarkers by the FDA [13]. Indeed, a recent comprehensive analysis of 6,729 pan-cancer tumors could only identify actionable mutations with therapeutic options available in clinical practice (FDA-approved or international guidelines) or reported in late phase (III–IV) clinical trials in 5.2% and 3.5% of the samples, respectively [14]. These figures coincide with clinical trial enrolment rates [1], where only 89 out of 1,640 of patients could enter genotype-matched treatment trials, the vast majority of which involved mutations in four genes, namely *PIK3CA*, *KRAS*, *BRAF* and *EGFR*. This highlights an acute need to expand the current repertoire of response biomarkers to cover more drugs and patients.

The eligibility criteria of most genomically-matched basket clinical trials are based on the single-gene biomarkers. However, most tumors do not present a single actionable mutation but have co-occurring driver alterations that might simultaneously alter key players of signaling pathways connected by cross-talk and feedback mechanisms [15, 16]. There are many documented cases of functionally relevant co-occurring oncogenic mutations, such as the concomitant inactivation of *TP53* and *RB1* [17], co-deletion of *CDKN2A* and *CDKN2B [18]*, co-amplification of *MDM2* and *CDK4* [19, 20], 1p/19q co-deletion in glioma [21], *MYC* amplification and *TP53* mutations [22] or activating alterations in *KRAS* and *BRAF* [5]. Indeed, although strong oncogenic KRAS and BRAF alterations are mutually-exclusive in treatment naïve tumors, a deeper allele-specific analysis identified a significant co-occurrence of activating RAS alterations and BRAF^D594^ mutations, among other co-occurring hotspot mutations within MAPK signaling genes (i.e MAP2K1 and upstream activating mutations in BRAF or NRAS) [16]. Additionally, at pathway level, the concomitant activation of PI3K signaling pathway with FGF signaling (*FGFR2* and *FGFR3*), or with *NRF2* mediated oxidative response have also been identified in several tumor types [16]. In this context, a single-gene based stratification of patients into subtypes and treatment arms might be over-simplistic, and novel frameworks that exploit co-mutational patterns might prove more effective.

As in the identification of driver mutations, the discovery of drug response biomarkers requires large numbers of patient molecular profiles matched to treatment outcomes. Unfortunately, treatment history information of large-scale genomics endeavors has not been systematically collected (e.g. TCGA [23]) or is not yet publicly available (e.g. GENIE Consortium [24]). Even though better data sharing policies are needed, many concerns are raised regarding privacy, property and the preliminary nature of confidential biomedical data. Safer alternative ways of sharing biomedical data are already on the table [25] but, until the access to systematically annotated clinical records becomes a reality, the research community largely relies on drug response data gathered from pre-clinical models.

Cancer cell lines are the most widely used *in vitro* model system, and have been fundamental tools to set the grounds of our understanding of cancer biology and to assess the efficacy of a broad spectrum of cancer drugs [26]. Unfortunately, cancer cell lines have been cultured as monolayers on plastic surfaces, and in growth-promoting conditions, for decades. As a consequence, most of them have suffered a substantial transcriptional drift, and they likely represent a cell subpopulation from the original primary tumor [27]. Those facts have fueled the debate regarding how well cancer cell lines resemble the tumors from which they were established and to which extent they are clinically relevant [15, 27]. A more realistic model to bridge the bench-to-bedside gap is the patient-derived mouse xenograft (PDX) [28]. To some extent, PDXs preserve inter- and intra-tumoral heterogeneity, and mimic the clinical course of the disease and response to targeted therapy, at least in certain tumor types [29–31]. Indeed, a recent review reported a 91% (153 out of 167) correspondence between the clinical responses of patients and their cognate PDXs [32]. Although this data are more time-intensive and expensive to generate, it is still feasible to establish large *in vivo* screenings, covering a wide diversity of tumor types and drugs. PDXs are thus a clinically relevant platform for pre-clinical pharmacogenomic studies, and represent a more accurate approach to identify predictive biomarkers compared with the use of cancer cell lines [33].

Here, we present a computational strategy to uncover and exploit driver alteration co-occurrence patterns in PDXs. By comparing the molecular profiles of responder and non-responder PDXs to a given drug, we identify driver co-occurrence networks and use them as a new type of drug-response indicator, applicable much beyond known biomarkers. We apply our strategy to the largest panel of PDXs and drugs available to date [28], and prospectively validate our findings *in vivo*. Finally, we adapt our strategy to derive response predictive models directly from continuous clinical outcome measures, such as progression free survival, and evaluate them on a cohort of breast cancer patients.

## Methods

### Genomic data processing

A total of 1,075 PDX models were established as part of a large pharmacogenomics screening that used the ‘one animal per model per treatment’ (1×1×1) experimental design to assess the population responses to 62 treatments [28]. We collected somatic mutations and copy number alterations for 375 of them, and used the *Cancer Genome Interpreter* resource [14] to classify protein-coding somatic mutations and copy number variants into predicted passenger or known/predicted oncogenic. In order to increase the clinical translatability, we subsampled both datasets to consider those oncogenic alterations covered by MSK-IMPACT [34] or by Foundation Medicine [35] targeted gene panels to obtain DCO networks and TCT4U models that could be directly used with those kind of molecular profiles, which are becoming widely used in the clinical setting.

### Drug response data

In the original dataset, a total of 62 treatment groups were tested in 277 PDXs across six indications. Drug response was determined by analyzing the change in tumor volume with respect to the baseline along time. They combined two metrics (Best Response and Best Average Response) into a modified RECIST classification (mRECIST) with four classes: PD (progressive disease), SD (stable disease), PR (partial response) and CR (complete response). For our analyses, we considered PDXs whose tumors progressed upon treatment (PD) as non-responders, and PDXs whose tumors stopped growing (SD) or regressed (PR, CR) as responders. After applying this binary classification, we had to exclude 9 treatments for which there were less than 5 PDXs in one of the two response groups, lacking thus enough interindividual heterogeneity to model drug response. A total of 276 PDXs were treated in at least one of the 53 treatment groups considered, each treatment being tested in 29 to 246 animals, with a median of 43 (IQR: 38-93). We could obtain the molecular profile for 187 of them, which had been treated with a median of 18 (IQR: 14-20) drugs. The final dataset consisted on 3,127 experiments performed on 187 PDXs and 53 treatment responses, across 5 tumor types: BRCA (breast cancer, n=38), CM (cutaneous melanoma, n=32), COREAD (colorectal carcinoma, n=51), NSCLC (non-small-cell lung carcinoma, n=27), PAAD (pancreatic adenocarcinoma, n=38), and 1 PDX without tumor type annotation).

### Molecular representativity of PDXs

We used the *OncoGenomic Landscapes* tool [36] to obtain a 2D representation of the molecular heterogeneity of the 187 PDXs being analyzed, and compared it to that of large reference cohorts of cancer patients. We downloaded the precomputed 2D projections of the following reference cohorts from the *OncoGenomic Landscapes* webserver (oglandscapes.irbbarcelona.org): PanCancer (n=15,212), BRCA (breast cancer, n=2,021), CM (cutaneous melanoma, n = 492), COREAD (colorectal carcinoma, n = 1,442), LUAD (lung adenocarcinoma, n = 1,486), LUSC (lung squamous cell carcinoma, n = 352), and PAAD (pancreatic adenocarcinoma, n = 442). We merged LUSC and LUAD samples in order to get a reference cohort for NSCLC (non-small-cell lung carcinoma) PDXs. We selected the 2D coordinates of the subset of TCGA and MSKCC patients of each reference cohort and represented their distribution in the PanCancer landscape as a level plot using the 2D kernel density estimate function of the ‘seaborn’ python library with 20 levels and a color-map that represents probability density as heat in the background. We selected the 2D coordinates of the 187 PDXs and represented their individual location with points, colored by tumor type.

### Drug response prediction based on Cancer bioMarkers database

We manually mapped the set of 53 drugs and drug combinations tested in the cohort of PDXs to the corresponding drug families in the *Cancer bioMarkers database [14]* using drug target information available in ChEMBL and DrugBank (Additional File 1: Table S1). We successfully assigned 50 out of the 53 treatments, spanning 29 drug family annotations. We considered those genomic alterations showing a ‘complete match’ with any of the reported predictive biomarkers and collapsed them at gene level. Note that we adopted a tissue agnostic approach in the development of TCT4U and, therefore, we did not require that the tissue of origin of the PDX matched the tissue or lineage in which each biomarker was identified. Nevertheless, this information is provided in Additional File 1: Table S2 to enable other researchers to perform stratified analyses. We considered as ‘approved’ biomarkers those ones that are currently approved by the FDA or by the main clinical guidelines in the field, such as the National Comprehensive Cancer Network (NCCN), the College of American Pathologists (CAP), the Clinical Pharmacogenetics Implementation Consortium (CPIC), or the European LeukemiaNet guidelines. We considered the rest of biomarkers, with varying supporting evidence, as ‘experimental’ biomarkers. The *Cancer bioMarkers database* usually reports more than one biomarker per drug or drug family, and often a single patient (or PDX) harbors several biomarkers of response and/or non-response for the same drug or drug family. We grouped response and non-response biomarkers at gene level and calculated the balanced accuracy (*BAcc*; average between sensitivity and specificity) of the prediction made by each gene in each treatment arm.

We weighted the binary predictions made by each gene and combined them to obtain a final prediction per treatment and PDX (*wComb*_*bmk*_).

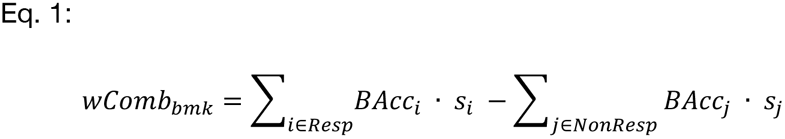

*Resp*: Set of genes with or without predictive biomarkers of response (*s*_*i*_, binary)

*NonResp*: Set of genes with or without predictive biomarkers of non-response (*s*_*j*_, binary)

*BAcc*: balanced accuracy of the predictive biomarker in a given treatment arm.

### Driver Co-Occurrence (DCO) Networks

#### Differentially altered drivers (DiffD)

For each treatment, we aimed at identifying differentially altered driver genes (DiffD) across responder and non-responder PDXs. To this end, we used *methyl_diff* [37], an analytical solution to estimate the probability of the inequality between the mutation rate of each driver gene across response groups, modeled using Beta distributions. We identified three sets of genes per treatment arm: (i) Resp_DiffD are those genes with more than 95% probability of showing higher alteration rate in responder PDXs than in non-responders, (ii) NonResp_DiffD are those genes with more than 95% probability of showing higher alteration rate in the non-responder than in responder PDXs, and (iii) General_DiffD are those genes with more than 95% probability of showing differential alteration rate between the two response groups. Additionally, we required that the selected genes were altered more than once in the corresponding group, with a minimum alteration rate of 5%.

#### Driver Pairs (Ps)

To identify pairs of driver gene alterations occurring more often than expected in each response group of a given treatment arm, we compared the observed co-occurrence rate to the random expectation under a null model with preserved sample- and gene-wise alteration rates. We obtained this null model by generating 1,000 random permutations of the genomic alteration matrix with the R package BiRewire [38]. We computed the average probability that the co-alteration rate observed in the actual dataset is larger than the co-alteration rate observed in the permuted datasets with *methyl_diff* [37]. When the average probability was larger than 95% we considered that the pair of drivers showed a tendency towards co-ocurrence.

Additionally, we computed the probability of the differential co-occurrence rate between responder and non-responder PDXs. We computed the same probability under the null model and compared its distribution to the probability of differential co-occurrence observed in the actual dataset. We selected the following sets of pairs per treatment arm: (i) Resp_Ps are those pairs showing a significant tendency towards co-occurrence in responder PDXs and showing a 95% probability of being co-altered more often in responder than in non-responder PDXs; (ii) NonResp_Ps, which show a significant tendency towards co-occurrence in non-responder PDXs and a 95% probability of being co-altered more often in non-responder than in responder PDXs; and (iii) General_Ps, which show a significant tendency towards co-occurrence in the whole treatment arm. Additionally, we required that the selected pairs were altered more than once in the corresponding group, with a minimum inferred alteration rate of 5%. In the case of Resp_Ps and NonResp_Ps, we additionally required that the probability of differential co-occurrence rate was larger than the 95% percentile of the distribution of probabilities obtained when comparing permuted samples.

#### Driver Co-Occurrence (DCO) networks

The differentially-altered drivers (General_DiffD, Resp_DiffD, NonResp_DiffD) and pairs of co-altered drivers (General_Ps, Resp_Ps and NonResp_Ps) can be expressed in terms of co-occurrence networks, in which nodes representing differentially altered driver genes (DiffD) or driver genes involved in a pair of co-altered drivers (DiP) are connected according to significant co-occurrences (Ps). For each treatment arm, we obtained three of such networks: (i) a general network (General_DCO), (ii) a responders network (Resp_DCO), and (iii) a non-responders network (NonResp_DCO).

### TCT4U drug response classifiers

We described the DCO networks with a matrix of Boolean vectors (1: altered, 0: unaltered) encoding the alteration status of differentially altered drivers and drivers participating in co-occurring pairs in each PDX (DiffD_DiP). We put together all those vectors in the form of a matrix and used it to train a decision-tree based gradient boosting classifier (CatBoost [39]). We did not specify the edges as features because CatBoost already considers first order interactions between all pairs of features, meaning that it natively exploits driver co-occurrences to predict treatment outcome. We used 100 trees with a maximum depth ranging from 1 to 7, a learning rate ranging from 0.2 to 1, and a coefficient at the L2 regularization term of the Logloss function ranging from 1 to 10. We chose the best set of hyperparameters based on the on the AUC obtained in the five-fold cross-validation of 30 iterations. Please, note that we repeated the same procedure for each treatment arm with each of the three DCO networks described before (General_DiffD_DiP, Resp_DiffD_DiP and NonResp_DiffD_DiP). We assessed the accuracy and robustness of each of the three classifiers by performing an external leave-one-out cross validation (LOOCV). We used the balanced accuracy of the LOOCV as weight to combine the General_DiffD_DiP, Resp_DiffD_DiP and NonResp_DiffD_DiP predictions generated for each drug-PDX instance into a final score, as described in Equation 2.

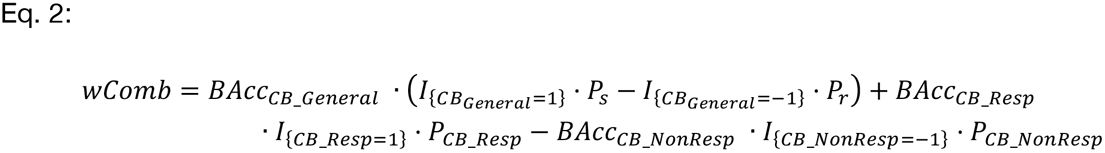

*CB*_*General*_, *CB*_*Resp*_, *CB*_*NonResp*_ : CatBoost binary predictions based on General_DCOs, Resp_DCOs, and NonResp_DCOs. Values of 1 and −1 indicate response and non-response.

*P*_*CB_General*_, *P*_*CB_Resp*_, *P*_*CB_NonResp*_: Probability estimates for the predictions.

*I*_*{CB=j}*_: Indicator function that takes a value of 1 when CatBoost predicts class j or a value of 0 otherwise.

*BAcc*_*CB*_: Balanced accuracy attained by a given CB classifier in the LOOCV.

We used exactly the same pipeline to obtain DCO-based predictions, biomarkers-based predictions and predictions based on an integrated model (DCO+biomarkers).

### Model interpretation

We computed the average feature importance in the LOOCV and excluded from the DCO networks those driver genes that did not contribute to the predictions. This effectively removed also a lot of pairs of co-occurring drivers that did not contribute either. We then quantified the feature’s local contribution to each individual prediction by analyzing the Shapley additive explanation (SHAP) values calculated within CatBoost.

Next, we assessed the specific contribution of co-occuring pairs of drivers to the prediction of drug response. To this end, we ranked driver pairs by the strength of the interaction, which is natively computed by CatBoost and used to generate drug response predictions. For each pair of drivers A and B, we classified the samples on the basis of the status of genes A and B. We then computed the average SHAP value of samples within each of the four resulting categories. We represented the results as SHAP interaction plots to uncover what is the effect of having a driver alteration in gene A on the SHAP value of gene B, and viceversa. We performed this analysis separately for pairs of drivers located in the same chromosome and pairs of drivers that are far apart in the genome.

### Experimental validation in PDXs

We collected all the available molecular profiles of the VHIO collection of breast cancer PDXs. Most PDXs were profiled using a hybridization-based capture panel of 410 genes (MSK-IMPACT) [34]. As we did for the training set, we used the Cancer Genome Interpreter resource [6] to filter out as many passenger alterations as possible. In the same way we did for the LOOCV, we described the molecular profile of each PDX according to the DiffD_DiP feature vectors associated to each DCO network and used them to predict the response to the 53 treatments in the TCT4U collection. For each PDX, we ranked all treatments based on the predicted response and focused on the predictions generated by the 21 treatments that attained a Balanced Accuracy of at least 0.6 in the leave-one-out cross validation. In order to increase the novelty of our findings, we prioritized those predictions that were not in agreement with predictions made by known predictive biomarkers. Of the drugs and PDXs available in our laboratories, we selected 8 positive and 6 negative predictions spanning the following treatments: MEK inhibitor (n=2), Pi3K inhibitor (n=5), taxane (n=2), Pi3K inhibitor + CDK4/6 inhibitor (n=3), and CDK4/6 inhibitor (n=2).

For each drug-PDX pair, 2 to 10 tumors were subcutaneously implanted in immunocompromised mice and grown until they reached a volume of 120-150 mm^3^. Tumors were treated with either vehicle or the corresponding drug or combination at a clinically relevant dose. Tumor growth was measured at least twice per week for approximately 20 to 40 days, when typically tumor volume in the control group had doubled twice or more. Caliper measurements were converted into tumor volume estimates using the formula (*l* · *w* · *w*) · (π/6), where *l* and *w* are the major and minor tumor axes, respectively. The response was determined following the mRECIST guidelines that were used in the PDX screening that we used as training set [28]. Basically, we calculated the percentage change in tumor volume from baseline (Δ*Vot*_*t*_ = (*V*_*t*_ − *V*_*i*_)/*V*_*i*_ · 100) and determined the BestResponse as the minimum value of *V*_*t*_ after 10 or more days of treatment. In order to capture tumor growth dynamics, we also calculated the BestAverageResponse as the minimum value of 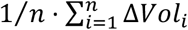 after 10 or more days of treatment. PDXs were classified into response groups according to the mRECIST criteria applied in the following order:

CR: BestResp < −95% and BestAvgResp < −40%
PR: BestResp < −50% and BestAvgResp < −20%
SD: BestResp < 35% and BestAvgResp < 30%
PD: BestResp ≥ 35% and BestAvgResp ≥ 30%

We used the R package Xeva [40] to generate the tumor response plots shown in Figure 5.

### Adaptation of TCT4U to use continuous clinical outcome measurements

We obtained both genomic and clinical data for a total of 216 patients with HR+/HER2− metastatic breast cancer that were treated with a CDK4/6 inhibitor in combination with an Aromatase Inhibitor in metastatic setting [41]. All patients underwent prospective clinical genomic profiling consisting on the identification of single nucleotide variants, small indels and copy number alterations detected from matched tumor-normal sequence data using the MSK-IMPACT targeted gene panels. We used the Cancer Genome Interpreter [14] to filter out passenger mutations and CNVs and keep only known or predicted driver mutations or copy number alterations. Detailed treatment history data was collected for each patient and included all lines of systemic therapy from the time of diagnosis of invasive carcinoma to the study data lock in September 2017. The exact regimen, as well as the dates of start and stop of therapy were also recorded. For the current analysis, we considered the treatment duration time as a measure of clinical benefit derived by patients whose biopsies were collected prior to or within the first 60 days of therapy initiation.

We used the TCT4U model of response to ribociclib to predicted response to CDK4/6 inhibition, as described before. Due to the differences in clinical outcome measurements between the training and the clinical cohort, we decided to adapt the TCT4U methodology to use continuous clinical outcome measurements as training set, instead of binary classification of drug response based on tumor growth. Our strategy consisted on comparing extreme populations both to derive the DCO networks and to train the classifier. We partitioned the population into three equally sized sets and applied the methodology described above. In this exercise, we set the cut-offs at 4.2 months and 9.7 months. We selected as Resp_DiffD or NonResp_DiffD those genes with more than 95% probability of showing higher alteration rate in the one third of patients showing the most durable or shortest clinical benefit, respectively, compared to the third of patients at the other extreme of the distribution. Additionally, we selected as General_DiffD all those genes with more than 95% probability of showing differential alteration rate between the two extreme populations. The same strategy was applied in the identification of pairs of driver gene alterations occurring more often than expected considering all patients (General_Ps) or separately for the one third of patients that relapsed the latest (Resp_Ps) or the earliest (NonResp_Ps). The remaining steps were applied exactly as described for the binary TCT4U methodology. In this setting with only one treatment per patient, high confidence predictions were selected by optimizing the threshold of the global score to get a maximum false discovery rate of 30% in the LOOCV, which happened when we kept predictions of response with a score above and 0.23 and predictions of non-response with a score below −0.26.

## Results

### Driver co-occurrence networks of drug response

Although thousands of genomic profiles of patient tumors are available, accurate information about pharmacological interventions and treatment outcome has not been systematically collected [23], or has not been disclosed yet [24]. Thus, to bypass these limitations, we compiled drug response data obtained in PDXs, since they preserve the overall molecular profile of the original tumor, and maintain its cellular and histological structure [32]. In particular, we based our study on 375 PDXs for which somatic mutations and copy number alterations have been acquired, together with their response to 62 treatments across six indications, using the ‘one animal per model per treatment (1×1×1)’ experimental design [28]. As suggested by the authors, we adopted the Modified Response Evaluation Criteria in Solid Tumors (mRECIST) [28, 42] to assess the change in tumor volume in response to treatment. We considered ‘responders’ those PDXs that showed a Complete Response (CR), Partial Response (PR) or Stable Disease (SD), and ‘non-responders’ those with a Progressive Disease (PD) status.

Of the 62 drugs and drug combinations tested, we selected 53 treatment arms that showed significant inter-individual heterogeneity (i.e. a sufficient number of ‘responder’ and ‘non-responder’ tumors) to model drug response. In total, these data comprised 3,127 experiments performed on 187 PDXs [28] for which we had, at least, 5 responder and 5 non-responder PDXs. First, we assessed whether this set of PDXs is representative of the genomic diversity observed in human tumors by comparing their alterations to the oncogenomic profiles extracted from 13,719 cancer patients [36]. We found that the 187 PDXs considered broadly covered the whole oncogenomic landscape represented by the full cohort (‘PanCancer’ cohort in Figure 1). When analyzing tumor types individually, we observed that, while the mutational diversity of some of them is perfectly reflected in the PDX samples (e.g. colorectal and cutaneous melanoma tumors), the distribution of mutated genes showed clear differences in others (e.g. NSCLC). As expected, we observe that PDXs sharing tissue-of-origin are more similar between them than to other PDXs and, more importantly, that the same level of similarity is maintained between PDXs and patient samples (Figure 1). Overall, there are PDXs representing the most populated areas of the PanCancer cohort, suggesting that the full collection of PDXs may be used in downstream analyses.

**Figure 1.**
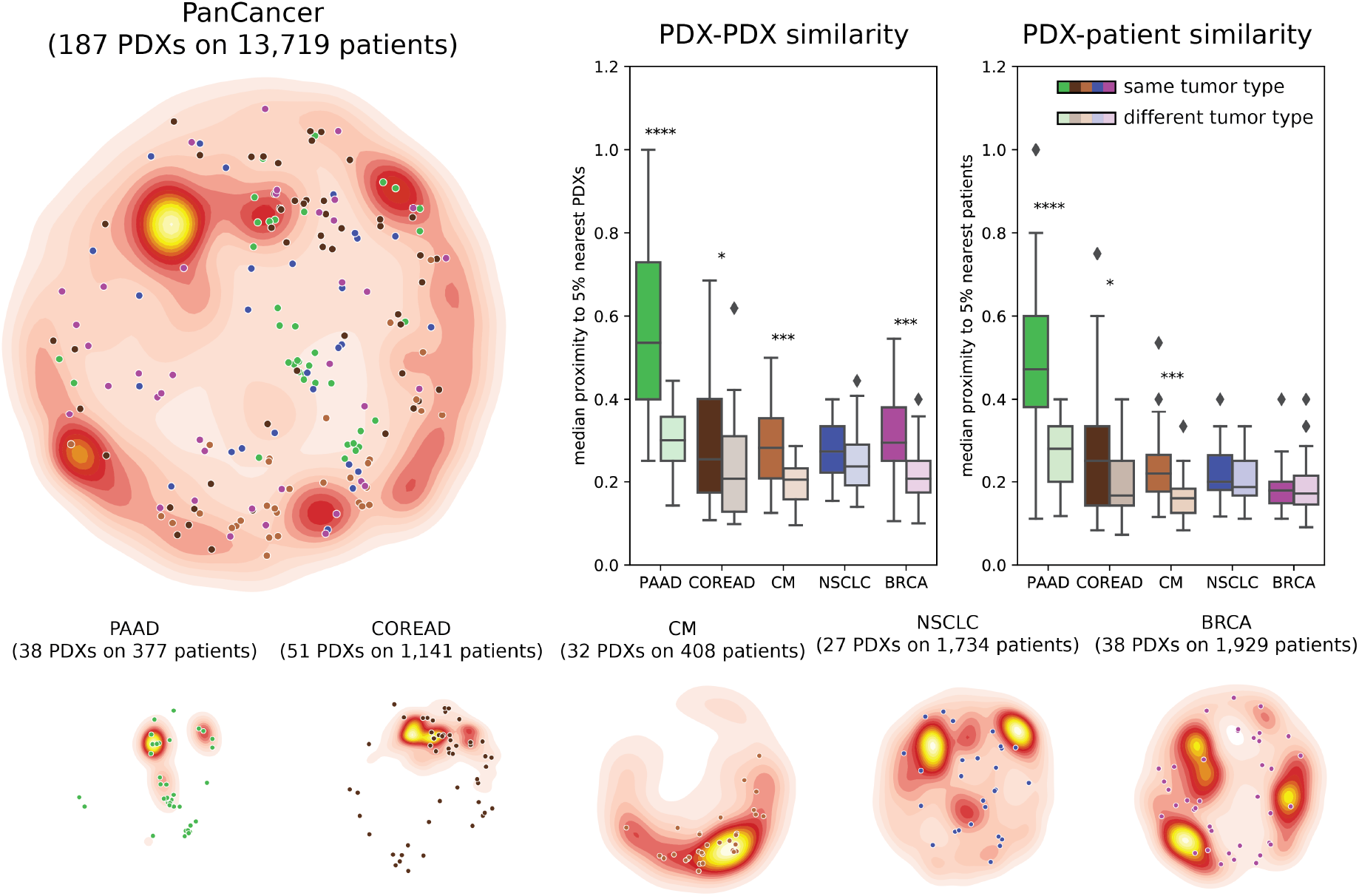
Molecular representativity of PDXs. OncoGenomic Landscape 2D representations of the molecular heterogeneity of the 187 PDXs annotated with both drug response data and oncogenic alterations, compared with that of their corresponding reference cohorts of cancer patients from TCGA and MSKCC. The points represent the location of each individual PDX, colored by tumor type. The distribution of the 187 PDXs can be compared to the distribution of patient samples, represented as density color-scale map in the background: PanCancer, PAAD (pancreatic adenocarcinoma), COREAD (colorectal carcinoma), CM (cutaneous melanoma), NSCLC (non-small-cell lung cancer), and BRCA (breast cancer). The boxplots show the proximity (median Jaccard similarity coefficient) of PDXs to the 5% nearest neighbors in each comparison. On the left, we show the clustering of PDXs based on tissue-of-origin by comparing the proximity of PDXs of a given tumor type among themselves and to PDXs of other tumor types. On the right, we show the clustering of PDXs with patient samples of the same tumor type compared to patient samples of other tumor types. Stars denote the p-value of a Wilcoxon rank-sum test (* <0.05, ** <0.01, *** <0.001, and **** p-value <0.0001).

We used the Cancer Genome Interpreter [14] to filter out passenger mutations from PDX profiles, and only worked with driver somatic mutations and copy number alterations. For each treatment, we grouped responder and non-responder PDXs, irrespective of the origin of their tumors. We first identified driver alterations that showed a differential mutation rate between responder and non-responder PDXs. Next, we identified pairs of driver alterations occurring more often than expected in each subpopulation under a null model that preserves both gene-wise and sample-wise alteration rates. Moreover, we identified pairs of driver alterations that show a differential co-alteration rate between responder and non-responder PDXs. With the overrepresented drivers (nodes) and pairs of co-occurring drivers (edges) identified, we built a Driver Co-Occurrence (DCO) network that characterizes responder PDXs, another DCO network that characterizes non-responder PDXs, and a third general one consisting of all drivers and co-occurrences associated with both treatment responses (Figure 2A).

**Figure 2.**
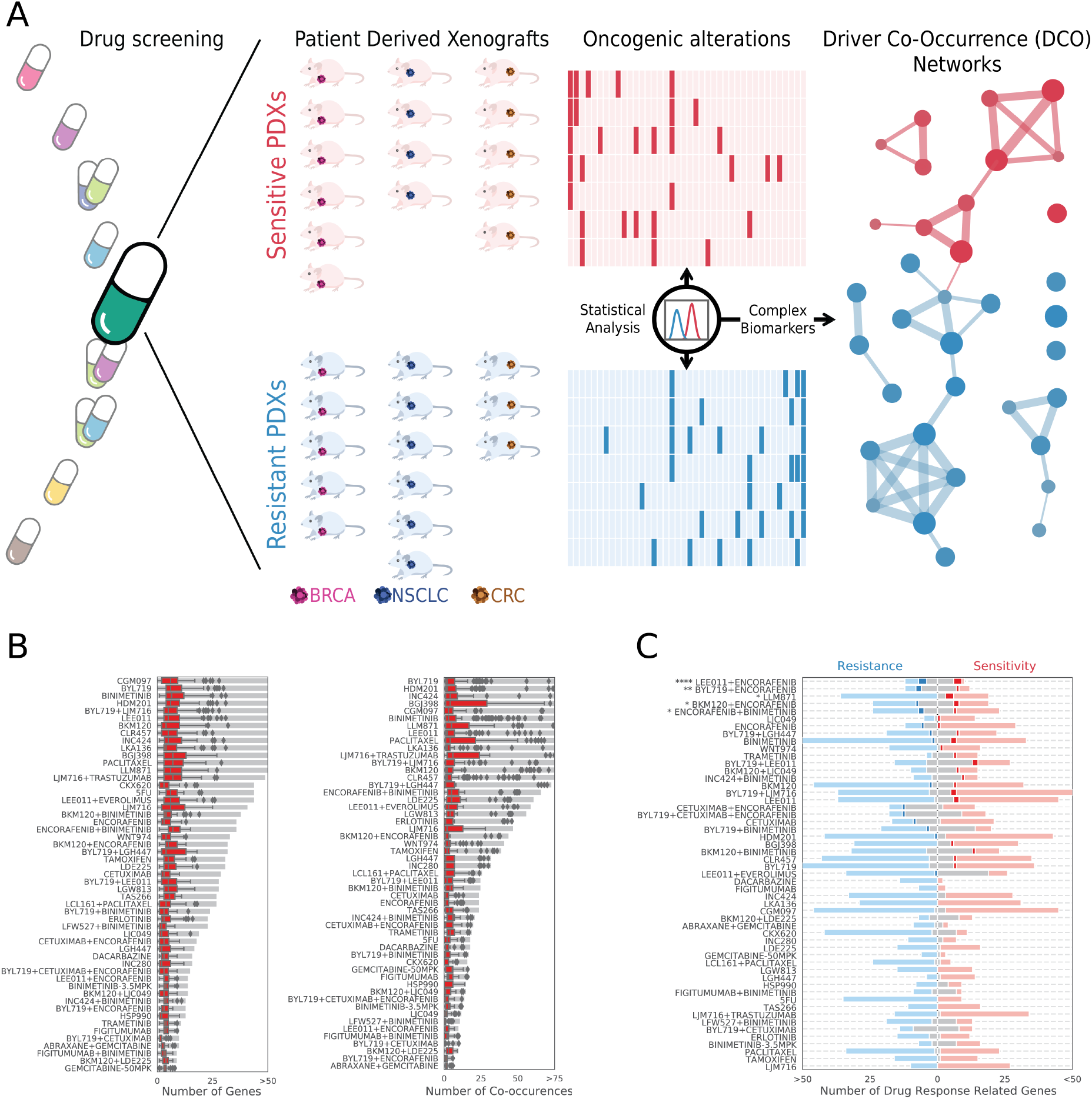
Computational strategy and description of Driver Co-Occurrence (DCO) networks. **(A)** We inferred DCO networks from the analysis of 3,127 *in vivo* experiments that screened the efficacy of 53 treatments against a panel of 187 molecularly characterized PDXs of several tumor types. We first compared the patterns of oncogenic mutations and CNVs in responder and non-responder PDXs, regardless of the tissue of origin of the tumors. Next, we identified sets of driver genes showing differential alteration rates between responders and non-responders (DiffD), which are represented as red or blue nodes in DCO networks, respectively. Additionally, we identified pairs of genes whose alteration co-occurred more often than expected under a null model with preserved gene-wise and sample-wise mutational frequencies (Ps), and that did so more often in one of the two response groups. We represented each pair of co-altered drivers as two nodes connected by an edge. We derived a responder, non-responder, and global DCO network for each treatment. **(B)** Gray bars show the number of drivers and pairs of co-occurring drivers included in each DCO network derived from whole exome sequencing data. Red boxplots represent the number of drivers or driver co-occurrences identified in each individual PDX. **(C)** Red and blue boxes represent the overlap between DCO drivers and genes with annotated biomarkers of response or non-response, respectively. We show in light colors the number of drivers in the DCO networks that were not previously associated to drug response. Likewise, gray bars indicate the number of previously known biomarker genes that were not included in our DCO networks. In this analysis, we only considered as eligible biomarkers those that were identified in two or more PDXs, which is a requirement that any driver needs to satisfy in order to be incorporated to a DCO network. Stars denote the p-value of a Fisher’s exact test (* <0.05, ** <0.01, *** <0.001, and **** p-value <0.0001).

DCO networks for each of the 53 drugs are detailed in Additional File 1: Table S2 and can be visualized using Cytoscape [43] (Additional File 2; https://doi.org/10.6084/m9.figshare.12789068.v1). The total number of drivers and driver co-occurrences captured in the DCO networks varied substantially among treatments, ranging from 8 to 89 driver genes (median of 29 nodes, IQR: 15-49) and 6 to 177 pairs of drivers (median of 26 edges, IQR: 16-93) overrepresented in PDXs treated with abraxane+gemcitabine and alpelisib (BYL719), respectively. However, when considering individual animals, the number of altered drivers and pairs of drivers was small and remained quite stable across treatments, with a median of only 4 genes (IQR: 3-5) and 2 driver co-occurrences (IQR: 1-3) per PDX (Figure 2B, Additional File 3: Figure S1).

We next sought to assess the novelty of our DCO networks by quantifying their overlap with the set of annotated response/non-response biomarkers for each treatment [14] that fulfill the eligibility criteria applied to build our DCO networks. Figure 2C shows that, although there is some overlap, our approach vastly expands the set of genes to be considered in downstream treatment prioritization applications. In some treatments we observe a significant overlap between DCO networks and known biomarkers (i.e LEE011+ENCORAFENIB, BYL719+ENCORAFENIB, or the FGFR inhibitor LLM871, Additional File 1: Table S3), but in most treatment arms the counts are too low to attain statistical significance. However, when aggregated across treatments, the overall overlap is significant, with 48 out of 359 possible drug-driver associations captured among the 1,856 total drug-driver pairs covered in TCT4U, from the universe of 30,687 eligible pairs (Fisher’s test OR 2.45, p-value 1.91·10^−7^, see Additional File 1: Table S3). Although we do not expect to recapitulate all previously reported drug-driver associations, finding some of them is a good sign of functional relatedness between DCO features and known mechanisms of action.

We are aware that without having performed a stratified analysis, we cannot rule out the possibility that tumor lineage might be a source of indirect associations between some genomic features and response to treatment, and we might miss interesting context specific biomarkers. However, we believe that, whenever identified, those biomarkers that are less context sensitive and that are common across different tumor types would be of special interest due to their wider applicability domain.

### TCT4U: A collection of 53 drug response classifiers for genome-driven treatment prioritization

We then explored whether the sets of differentially (co-)altered genes in responder and non-responder PDXs can be used to predict treatment outcome. For each drug, we used the DCO networks to statistically classify PDXs as non-responder or responder. The goal of this exercise is to identify, among the available treatments, the best possible option for each individual based on its oncogenomic profile. We thus named the set of developed drug response classifiers Targeted Cancer Therapy for You (TCT4U).

In brief, for each treatment arm, we combined the probabilities assigned by three gradient boosting classifiers (CatBoost), trained with response, non-response and general DCO networks, into a single prediction score per drug-PDX pair (Figure 3A). To increase the clinical translatability of our approach, we repeated the calculations considering only those alterations detectable by the Memorial Sloan Kettering-Integrated Mutation Profiling of Actionable Cancer Targets (MSK-IMPACT) [34, 44] and the Foundation Medicine (FM) gene panels [35], which contain probes to detect 410 and 287 mutated genes, respectively, and are widely used in clinical settings. Finally, we assessed the effectiveness of TCT4U by comparing its predictive power to that of FDA-approved and experimental biomarkers (see *Materials and Methods* for details).

**Figure 3.**
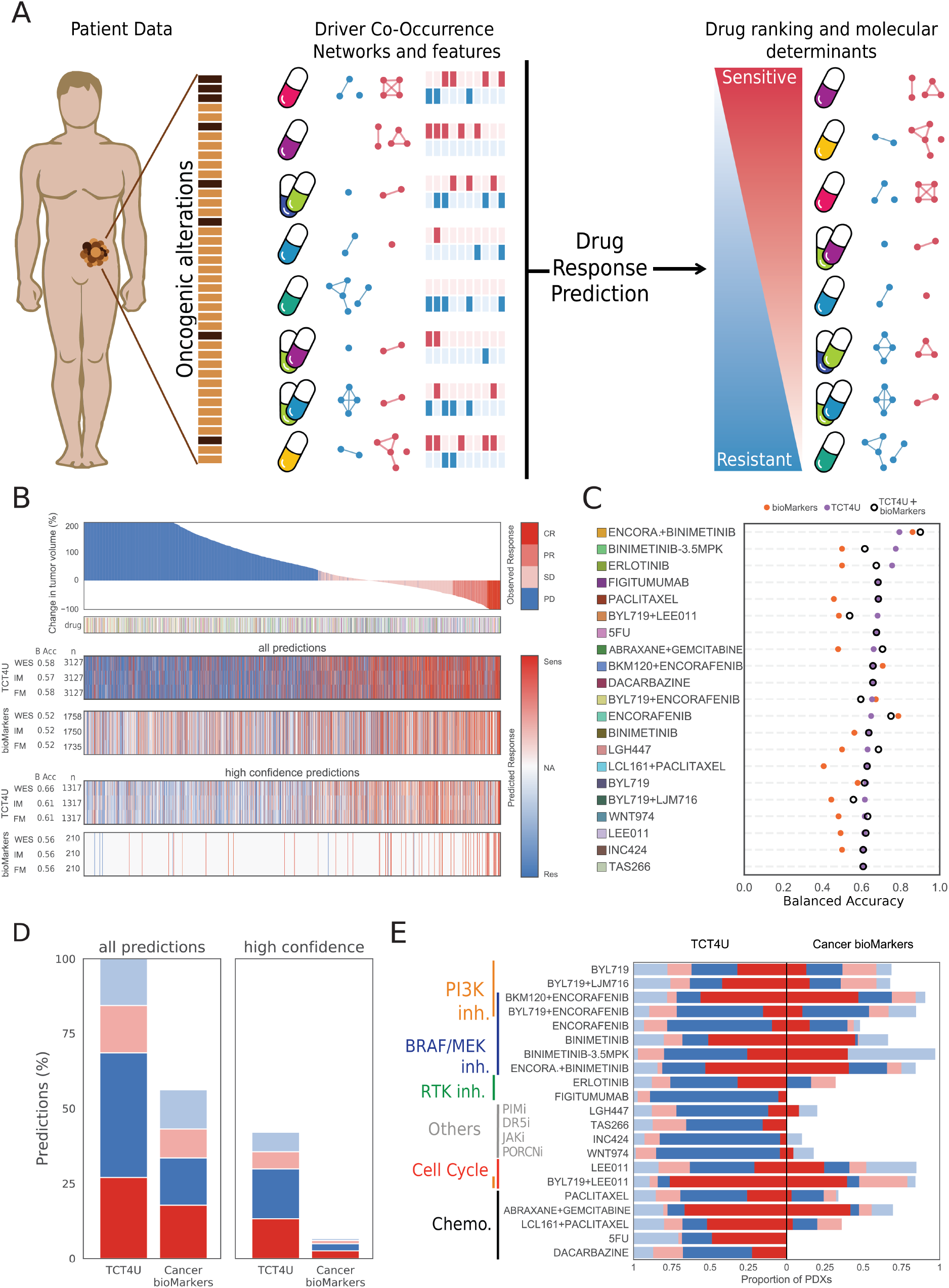
Targeted Cancer Therapy for You (TCT4U), a collection of drug response classifiers based on DCO networks. **(A)** Given a new tumor sample, we compare it to the patterns of driver alterations and co-alterations associated to response or lack of response to any of the treatments in TCT4U, and rank the drugs accordingly, predicting whether a drug will or not be effective. By quantifying each feature’s local contribution to individual predictions (SHAP values) we can know which are the precise molecular determinants used by the classifier and use this information for functional interpretation of the predictions. **(B)** Waterfall plot representing 3,127 *in vivo* experiments, sorted left to right from the worst to the best response of 187 PDXs to 53 treatments. The heatmaps below show the predictions of TCT4U in a leave-one-out cross validation setting, and the predictions made on the basis of approved or experimental biomarkers. Each heatmap has three rows, which correspond to the predictions obtained when examining the whole exome (WES), IMPACT410 (IM) or Foundation Medicine (FM) targeted gene panels. The number of predictions and the average balanced accuracy (BAcc) across treatments are annotated along the y-axis. The subset of high confidence TCT4U predictions corresponds to the ones yielded by models that attained a BAcc of 0.6 or higher. The subset of high confidence biomarkers corresponds to the subset of clinically approved ones. **(C)** Predictive performance of TCT4U, biomarkers and DCO+biomarkers based models, all trained with CatBoost within the TCT4U pipeline. **(D)** The precision of each set of predictions is illustrated by the red and blue sections of the stacked bar plots, which represent the proportion of correct response and non-response predictions. Analogously, incorrect predictions are represented in faint colors. Missing predictions (NA) are represented in white to offer a comparative overview of the recall. **(E)** Stacked bar plots representing the precision and recall of all TCT4U predictions and all previously known biomarkers covered by WES profiles split by treatment arm.

We collected the change in tumor volume and the mRECIST classification for a total of 3,127 experiments with reported treatment outcome, comprising 187 PDXs tested for response to 53 treatments. Figure 3B shows the predictive performance of the models in a leave-one-out cross-validation setting, whereby the genomic profile of PDXs is used to predict response to each treatment. We observe that TCT4U models are applicable to all drug-PDX pairs (3,127), while alterations in approved and experimental biomarkers can only be found in about half of them (1,758). However, wherever applicable, both methods attain a similar overall accuracy. As many treatment arms do not have a balanced number of responders and non-responders, we quantified the balanced accuracy separately for each treatment and calculated their average (Figure 3C, Additional File 1: Table S4). TCT4U and known biomarkers attained an average balanced accuracy of 0.58 and 0.52, respectively. The balanced accuracy was greater than 0.6 for 21 treatment arms, yielding a total of 1,317 high-confidence predictions. Although this subset of predictions only covered 42% of all drug-PDX pairs, their average balanced accuracy improved to 0.66 (Figure 3C, Additional File 1: Table S4).

In the case of known and experimental biomarkers, they could only predict with a balanced accuracy higher than 0.6 the response to 4 of those 21 treatments, namely encorafenib, encorafenib+binimetinib, encorafenib+alpelisib (BKM120), and paclitaxel. As for FDA approved biomarkers, they predicted drug response in 210 of the drug-PDX pairs, spanning 59 PDXs and 13 treatments. Due to sample imbalance, the balanced accuracy could only be calculated for a subset of treatments, making an average of 0.56 (Figure 3D, Additional File 1: Table S4). It is also remarkable that, even if they consider a much lower number of genes, both MSK-IMPACT and FM derived models achieved comparable prediction accuracies (Figure 3B, Additional File 1: Table S4).

The comparison of TCT4U to the simple combination of biomarkers showed an overall improvement with respect to the current standard. However, given the way we modeled the treatment decision setting, not having a given biomarker does not contribute to the predictions (i.e not having NRAS alteration does not predict for response to BRAF inhibition). In order to maximize the information extracted from previously reported biomarkers and perform a more controlled comparison, we decided to use exactly the same pipeline implemented for TCT4U but training the classifiers with the few features representing known biomarkers. Reassuringly, the resulting predictions were strongly correlated with the predictions based on the simple combination of biomarkers (Spearman’s rho 0.32, p-value 1.2·10^−100^, Additional File 3: Figure S2A). Known biomarkers were only able to predict response to 6 treatments with a balanced accuracy greater than 0.60, even when using gradient boosting classifiers instead of their simple combination. It is noteworthy that the six treatments consist on encorafenib used alone or in combination with alpelisib, buparlisib, binimetinib and cetuximab. Figure 3C shows that known biomarkers outperform TCT4U in the only four treatments involving encorabenib (256 predictions in total), while biomarker based models did not perform much better than random in the remaining 17 treatments (Additional File 1: Table S4).

Further, we built integrated models containing both DCO-based features and known biomarkers, which slightly outperformed both DCO-only and biomarkers-only models, suggesting that the two sets of features carry orthogonal information (Figure 3C). However, the overall performance taking into account all treatment arms did not change much with respect to the DCO-based models (Additional File 1: Table S4, Additional File 3: Figure S2B-C).

Finally, while the coverage of approved or experimental biomarkers is mostly limited to BRAF/MEK inhibitors, PI3K/mTOR inhibitors or cell cycle related treatments, the predictions made by TCT4U also covered other drug families including chemotherapies, RTK inhibitors, and more experimental treatments targeting *Wnt signaling* (WNT974), or apoptosis related pathways (TAS266, LGW813) (Figure 3E).

### Interpretation of DCO-based predictions

In addition to the prediction performance of each model, it is key to evaluate their potential to uncover patterns that can generate new hypothesis and propose novel biomarkers. Current tree-based explanation methods allow us to understand how the model uses input features to make predictions. Beyond assigning the importance of each feature to the global prediction, we can compute the Shapley additive explanation (SHAP) values to quantify the local contribution of each feature to the individual predictions (i.e how important is each driver gene for predicting drug response in a given PDX).

As expected, in targeted therapies that are currently approved in biomarker specific indications, the SHAP values of their intended targets show that they usually contribute to DCO-based predictions. For instance, in most PDXs *BRAF* alteration predicts for response to binimetinib (MEK inhibitor) used alone or in combination with encorafenib (BRAF inhibitor). On the other hand, *KRAS* and *NRAS* alterations, which are upstream of *BRAF,* contribute positively when predicting response to binimetinib but negatively when predicting response to its combination with encorafenib (Figure 4A). Those are the kind of predictions that are successfully achieved by simple biomarkers (Figure 3C, Additional File 3: Figure S2). However, it becomes clear that, even when looking at an FDA approved biomarker such as *BRAF*, single driver gene alterations do not contribute equally in all PDXs because of the effect of their interaction with additional driver alterations.

**Figure 4.**
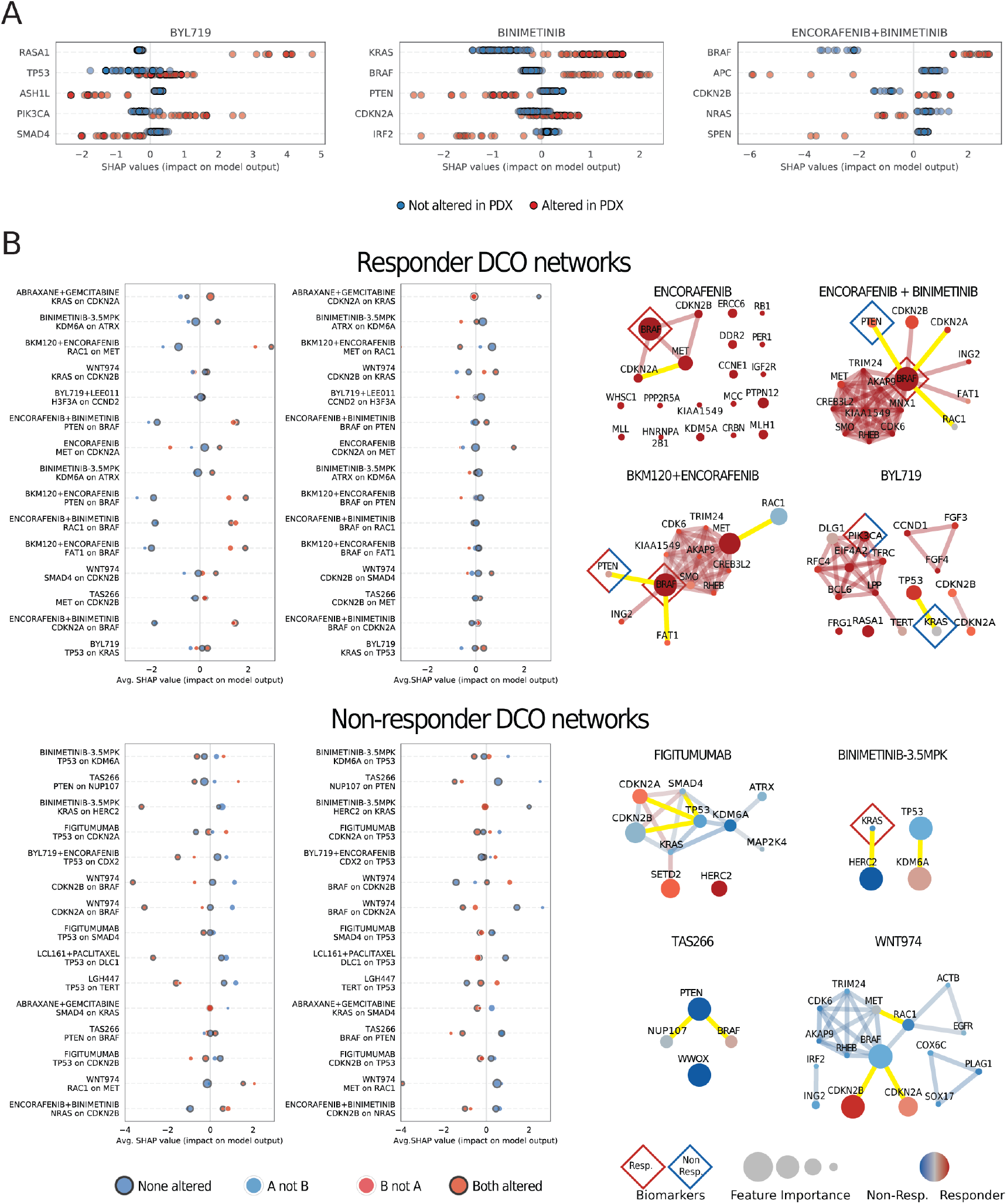
Specific contribution of driver co-occurences to the prediction of drug response. **(A)** Summary plot of the local feature’s contribution (SHAP values) attributed to individual genes when predicting response to treatment with TCT4U. Each point represents the contribution of a driver gene to the prediction of response in a given PDX. The color of the points indicate whether the given driver gene was altered or not in each PDX. The three examples represent the five most explanatory genes when predicting response to three approved targeted therapies with biomarker specific indications. **(B)** SHAP interaction plots and Driver Co-Occurrence networks (DCO) representing the oncogenic alterations and pairs of alterations that are overrepresented in responder and non-responder PDXs. SHAP interaction plots show the effect of having driver a alteration in gene A on the distribution of SHAP values of gene B. Each point represents the average SHAP value of PDXs classified on the basis of the status of the two drivers that tend to be co-altered. The size of the points is proportional to the number of PDXs that belong to each of the four resulting categories. The figure represents feature interactions involving driver genes that are located far apart in the genome. We also show some exemplary DCO networks with the driver co-occurrences represented in the accompanying SHAP interaction plots highlighted in yellow. The size of the nodes represents the average feature importance in the LOOCV, and their color represents the probability of being overrepresented in responder (red) or non-responder (blue) PDXs. Previously known biomarkers detected in the cohort are annotated with diamond shapes.

*PIK3CA*-mutant tumors are sensitive to isoform-selective PI3K inhibitors such as alpelisib (BYL719) [45–47]. Accordingly, we observed a higher response rate (65%, 15 out of 23) among PDXs with oncogenic *PIK3CA* alterations compared to PDXs with wild-type *PIK3CA* (44%, 52 out of 117), and the average SHAP value of *PIK3CA* in altered PDXs is positive (1.16, Additional File 1: Table S2). However, *PIK3CA*-independent mechanisms of PI3K activation (e.g. activating alterations in *PIK3CB* or *PTEN* loss) often limit the response to this treatment [48, 49]. The alpelisib DCO network contains three proteins involved in PI3K signaling, namely *PIK3CA*, *PIK3R1*, and *PIK3C2B.* Interestingly, we found that *PIK3CA*-altered PDXs having no co-occurring oncogenic alterations in the PI3K pathway showed an even higher response rate (79 %, 11 out of 14) than those with co-occurring alterations in *PIK3R1* or *PIK3C2B* (44%, 4 out of 9). This is reflected in the negative average SHAP values observed for *PIK3R1* and *PIK3C2B* in altered PDXs (−0.83 and −0.12, Additional File 1: Table S2). Our DCO-based model was able to capture alterations that are likely to activate PI3K signaling in a *PIK3CA*-independent manner, which could limit the response to alpelisib [49].

We extended this analysis to systematically assess the interaction between all co-occurring driver alterations in the 21 treatment arms that yielded high confidence predictions. We identified a total of 253 unique feature interactions between co-occurring drivers (Additional File 1: Table S2). Topological analysis of DCO networks revealed that several large, strongly connected modules were composed of genes that tend to be co-amplified or co-deleted as part of the same genomic segment. Although in most of the cases those clusters of co-altered drivers are probably hitchhiking with a major driver alteration that is positively selected, in some other cases the co-amplification and simultaneous overexpression of adjacent oncogenes can provide a basis for cellular cross-talk.

For example, the co-amplification of the two FGFR ligands FGF3 and FGF4 (chr11:q13) contributes positively to the prediction of response to alpelisib (average SHAP 1.59), while the alteration of FGF4 alone contributes negatively (average SHAP −1.34). Similarly, the co-alteration of MET (chr7:q31) and its downstream signal transducer BRAF (in chr7:q34, 24Mb away) contributes more negatively to predict response to the inhibition of Wnt signaling by WNT974 than any of the two alterations alone (Additional File 3: Figure S3B, Additional File 1: Table S2). In the same direction, MET alteration has also a negative impact on the prediction of response to encorafenib, encorafenib+buparlisib (BKM120), and encorafenib+binimetinib in BRAF altered PDXs (Additional File 3: Figure S3A, Additional File 1: Table S2). On the other hand, the same co-alteration contributes more positively to predict response to encorafenib and encorafenib+binimetinib than BRAF or MET alterations alone (Additional File 3: Figure S3A, Additional File 1: Table S2). Interestingly, a feedback loop between those two genes has been shown to influence response to BRAF and/or MET inhibitors [45].

Not surprisingly, the two aforementioned pairs of drivers are also co-altered more often than expected in a non-redundant set of 46,697 cancer patient samples queried through cBioPortal [50, 51] (log2OR 2.21 and >3, respectively; both p-values < 0.001). It has been suggested that the amplitude of the regions affected by copy number changes strongly determines patient prognosis [52, 53]. Broader amplifications are likely to modify the dosage of multiple genes which, based on our observations, could have a different impact on drug response than the focal alteration of a single driver gene.

However, when co-altered drivers are genomically linked it is very difficult to disentangle which is the specific contribution of each alteration because they would still co-occur even if only one of them was actually contributing to differential drug response. For this reason, we decided to distinguish between interactions involving genes located in the same chromosomal arm (Additional File 3: Figure S3) from those involving genes that are far apart in the genome (Figure 4B).

*As mentioned above, BRAF* driven tumors show a higher response rate (83%, 15 out of 18) when treated with encorafenib+buparlisib (BKM120) than *BRAF* wild-type tumors (36%, 5 out of 14), with an average SHAP value of 1.43 in altered vs −2.06 in non-altered PDXs. However, the SHAP value is even more positive for the subset of tumors with co-alteration of *PTEN* and *BRAF* (1.89). Indeed all tumors with this co-alteration responded to the treatment. On the contrary, *PTEN* deficiency in *BRAF* wild-type tumors is a negative predictor of response (average SHAP value of −2.60). Although to a lesser extent, we observed the same trend in the encorafenib+binimetinib model (Figure 4B). Importantly, *PTEN* and *BRAF* are also co-altered more often than expected in cancer patients queried through cBioPortal (OR 1.86 p-val < 0.001).

Another remarkable example is the interaction between *HERC2* and *KRAS* in the prediction of response to BINIMETINIB-3.5MPK (MEKi). This regimen was tested on 25 pancreatic tumors, most of which were *KRAS* driven (24 out of 25). Despite this fact, the downstream inhibition of *MEK1/2* was only effective in 62.5% of them (15 out of 24). Interestingly, *HERC2* was co-altered with *KRAS* in a substantial fraction of non-responder KRAS-driven tumors (25%, 5 out of 20). Accordingly, *HERC2-KRAS* co-alteration contributes negatively to the prediction of response (average SHAP value −3.22), whereas having wild-type *HERC2* and altered *KRAS* contributes positively (0.54). In patients, *HERC2* and *KRAS* are also significantly co-altered (cBioPortal OR 1.77, p-value < 0.001).

Overall, those example illustrate how TCT4U is able to detect and exploit interactions between driver alterations to predict drug response in PDXs. Sometimes these interactions involve previously known biomarkers, as illustrated by the co-alteration of BRAF and PTEN, which would have antagonistic effects in predicting response to the combined inhibition of BRAF and PI3K signaling. Moreover, many of these co-occurrence patterns are also found in patient cohorts, indicating a potential clinical translation of these findings. We provide the full set of interactions and their impact on SHAP values in Additional File 1: Table S2.

### Experimental validation of TCT4U drug response predictions on a prospective PDX dataset

In addition to the *in silico* benchmarks, we sought to prospectively evaluate the performance of the TCT4U models in new tumors. To this aim, we selected, from our VHIO collection of molecularly-characterized breast cancer PDXs, a subset of 14 drug-PDX pairs, with 8 tumors predicted to respond and 6 predicted not to respond namely: alpelisib, an isoform-selective PI3Kα inhibitor (BYL719, n=5); ribociclib, a CDK4/6 inhibitor (LEE011, n=2); the combination of both (alpelisib+ribociclib, n=3); the MEK inhibitor binimetinib (n=2); and paclitaxel, a taxane (n=2). This is a particularly challenging set of PDX-drug compounds since, except in one case, the anticipated TCT4U outcome did not agree with approved or experimental biomarkers, either because the individual genomic profiles did not have any biomarker altered (n=8), or because the TCT4U predictions were opposed to those suggested by known biomarkers (n=5).

We subcutaneously implanted the tumors in immunocompromised mice and let the tumors grow until they reached a volume of 120-150 mm^3^. We then treated the PDXs for 15-57 days and measured their response to the administered drugs following the mRECIST guidelines (see Materials and Methods for details). The complete results of our study, including TCT4U predictions, known biomarkers, treatment setting (drug dose, duration, etc.), and tumor response (tumor growth, mRECIST classification, etc.) for every PDX can be found in Additional File 1: Table S5, and are summarized in Figure 5.

**Figure 5.**
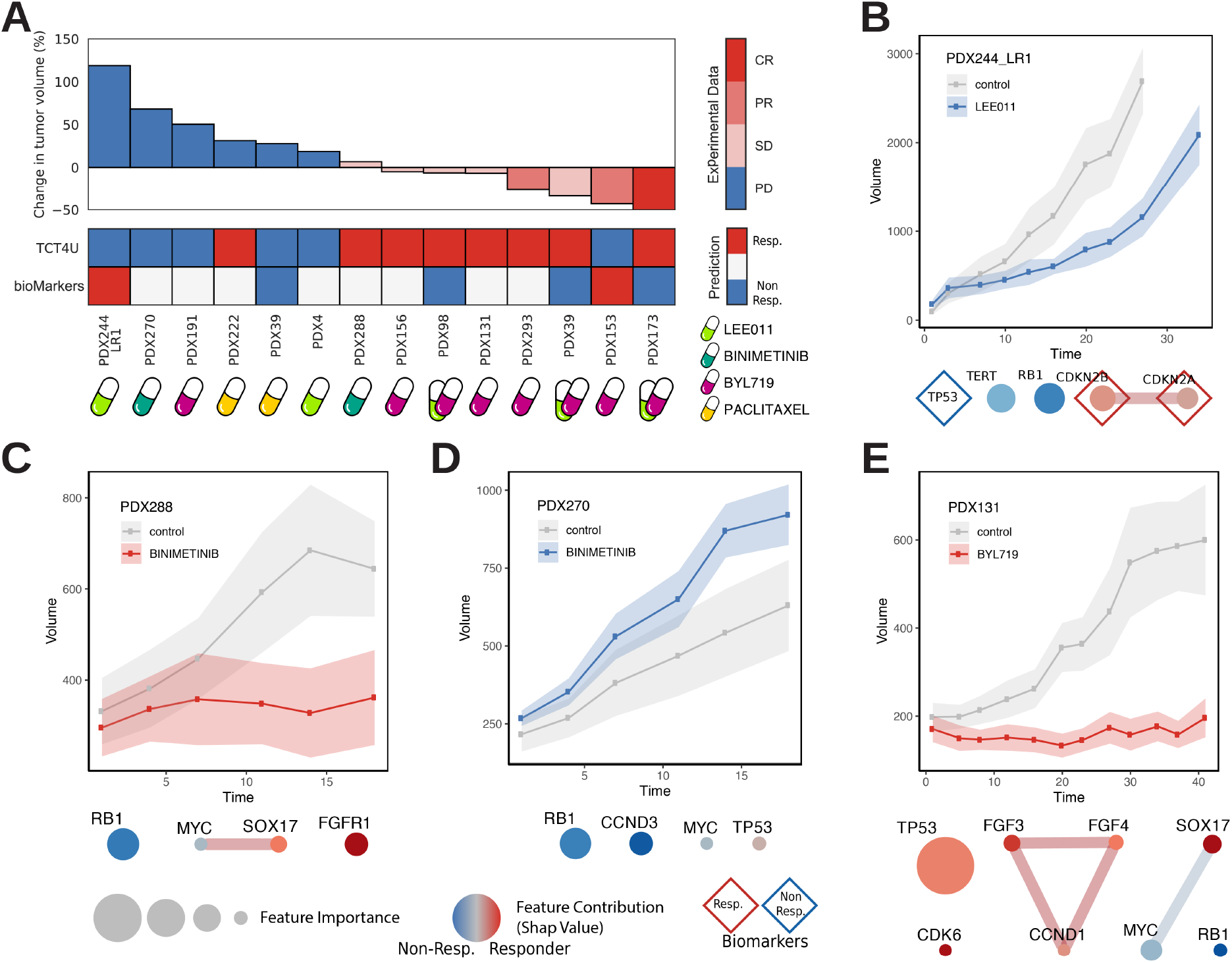
Experimental validation of TCT4U predictions. **(A)** The waterfall plot summarizes the results of 14 *in vivo* experiments comprising 5 PDXs treated with the alpelisib (BYL719) isoform-selective PI3K inhibitor (PI3Ki), 2 PDXs treated with ribociclib (LEE011) CDK4/6 inhibitor (CDKi), 3 PDXs treated with the combination of both (PI3Ki+CDKi), 2 PDXs treated with binimetinib MEK inhibitor (MEKi), and 2 PDXs treated with paclitaxel. The heatmap below shows the predictions based on TCT4U compared to previously known biomarkers. **(B-E)** Exemplary tumor response curves and features in the DCO network explaining TCT4U predictions. The size of the nodes represents the average feature importance in the LOOCV. The color of the nodes represents the contribution of each feature to the prediction of response in the given tumor (SHAP values). Previously known biomarkers detected in a given tumor are annotated with diamond shapes. The complete results of our study are provided in Additional File 1: Table S5.

We treated five PDXs with alpelisib, three of which (PDX131, PDX293, and PDX156) were predicted as responders to the drug by TCT4U models, and two as non-responders (PDX191 and PDX153). The three PDXs predicted as responders showed a co-amplification of the *FGF3-FGF4-CCND1* triplet, located in the 11q13 genomic segment. In our DCO-models, this triplet happened more often in responder than in non-responder PDXs, with an alteration rate of 7.46% and 1.37%, respectively. The three genes contributed positively to the prediction of response in these PDXs, with average SHAP values of 0.72, 1.86 and 0.26 (Additional File 1: Table S5). It is worth noting that our model, which was derived from 140 PDXs of different tumor types (i.e. 38 BRCA, 42 COADREAD, 25 NSCLC and 35 PDAC), did not show a significant tendency towards co-occurrence of PIK3CA and the 11q13 amplicon (OR 2.69 p-value 0.26). Dysregulation of FGFR signaling can lead to downstream activation of PI3K/AKT pathway and, indeed, a recent study reported that 73% of patients (8 of 11) with both an alteration in the PI3K/AKT/mTOR pathway and FGF/FGFR amplification experienced clinical benefit when treated with therapy targeting the PI3K/AKT/mTOR pathway, whereas only 34% of patients (12 of 35) with PI3K/AKT/mTOR alterations alone did so [54]. However, the implication of FGF signaling with respect to the clinical benefit of PI3K/AKT/mTOR blockage remains controversial. The retrospective analysis of a large subset of patients enrolled in the BOLERO-2 trial [55] showed that alterations in FGF signaling had a negligible impact (*FGFR1*) or slightly decreased (*FGFR2*) the clinical benefit of everolimus treatment. In line with these findings, ER+/*ERBB2-* metastatic breast cancer patients with *FGFR1* and *FGFR2* amplification did not derive a clinical benefit from alpelisib+letrozole [56]. Accumulating evidence suggests that FGF signaling induced by FGFR1/2 amplification attenuates the response to PI3K blockage in PIK3CA mutant breast cancer. However, the impact of FGF signaling in response to alpelisib in PIK3CA wild-type tumors originated from breast, as well as from other tissues, has yet to be determined.

In our dataset, the three PDXs responded to the treatment. In particular, in PDX293 we observed a partial response (PR) after 18 days of treatment, with a reduction of 65% in the initial tumor volume. PDX131 and PDX156 showed a stable disease (SD) after 20 and 11 days of treatment, respectively. On the other hand, PDX191 was predicted to be non-responder because, in addition to the aforementioned 11q13 amplicon, it had driver alterations in FGFR1, MYC, and GNAS that were contributing negatively to the prediction of response (SHAP values of −1.81, −1.04, and −0.73). In agreement with our prediction, the tumor increased its volume by 80% after 13 days of treatment (PD) (Additional File 1: Table S5). PDX153 was the only PDX with an oncogenic PIK3CA mutation (p.K111E) reported to confer sensitivity to the treatment [14] and, indeed, we observed a significant reduction of 83% in the tumor volume after 35 days of treatment (i.e. a PR outcome). Our model classified this PDX as non-responder because it also had other alterations overrepresented among non-responder PDXs, such as *MAP2K4* (SHAP value −2.85), or *NCOR1* (SHAP value −1.68). The DCO networks also considered *PIK3CA* status, which tends to be more frequently altered in responder PDXs (22.29%) than in non-responder PDXs (11.40%; SHAP value 1.86). However, the final prediction was driven by additional oncogenic alterations that together showed a stronger statistical association than *PIK3CA* status, although they proved to be less informative.

We administered ribociclib, a CDK4/6 inhibitor, to PDX4 and PDX244_LR1, with the TCT4U prediction that the two tumors would not respond to the drug. PDX4 did not present any known biomarker of drug response and the most influential feature in the ribociclib DCO-model was GNAS amplification. While the co-alteration of GNAS and AURKA is usually predictive of response to ribociclib (average SHAP value 0.461), the alteration of GNAS in the absence of additional alterations in this specific PDX contributed negatively to the prediction, with a SHAP value of −0.88 (Additional File 1: Table S5). On the other hand, we also treated PDX244_LR1, which is a model of acquired resistance to ribociclib derived from a responder parental tumor (PDX244). Accordingly, PDX244_LR1 simultaneously showed known biomarkers of response (*CDKN2A*-*CDKN2B* co-deletion) and non-response (*TP53* p.C176R) to the treatment [14]. *TP53* was not included in the ribociclib DCO network because we did not find it to be differentially altered between responders and non-responders (44.65% vs. 48.39%). In line with what has been reported, *CDKN2A*-*CDKN2B* co-deletion was slightly more common in responder than in non-responder PDXs (32.78% vs. 25.34) and contributed to positively to the prediction with SHAP values of 0.116 and 0.249 (Additional File 1: Table S5). However, PDX244_LR1 presents an oncogenic mutation in *RB1* (p.M695Nfs*26), which showed a strong association with lack of response to CDK4/6 inhibition in the DCO networks (4.29% vs. 12.65%) and is the most negative explainer in this model, with a SHAP value of −1.226. *RB1* is the primary target of CDK4/6 and its status is a key determinant of CDK4/6 inhibition efficacy [57]. Accordingly, *RB1* overexpression is reported to confer sensitivity to CDK4/6 inhibition in prostate cancer, although its loss or deletion is not currently reported as a non-response biomarker [14]. Our experiments showed that, and in agreement with TCT4U predictions, the tumors increased their volume between 45 and 215%, being thus catalogued as PD.

We also treated three PDXs (PDX173, PDX98 and PDX39) with the same PI3Kα and CDK4/6 inhibitors in combination (alpelisib+ribociclib). The three of them had oncogenic mutations in *TP53* (p.R249S, p.R249S and p.V157I), which are associated with non-response to CDK4/6 inhibition [14]. However, DCO networks found additional response-associated genomic features (i.e *MYC* in PDX98 with SHAP value of 0.558, or *H3F3A* in PDX39 with SHAP value of 0.321, Additional File 1: Table S5) and thus TCT4U models predicted them to respond to this drug combination. We found that, indeed, all three tumors responded to the combination treatment: PDX173 became completely tumor free (CR), PDX39 showed a reduction of 47% (SD), and PDX98 of 25% (SD).

Two PDXs were treated with the MEK inhibitor Binimetinib, with TCT4U models predicting PDX270 to be non-responder and PDX288 to respond to the drug. Both PDXs presented *RB1* loss (a loss-of-function mutation p.Y321* and a deletion, respectively), which is overrepresented in the non-responder DCO network (5.50% vs. 17.46%) and contributed negatively to the prediction (SHAP value of −0.549 in both cases). Additional alterations necessarily contributed to the divergent prediction of these PDXs. Alterations in *CCND3* and *MYC* contributed to the prediction of non-response in PDX270, with SHAP values of −2.147 and −0.330. *MYC* was also altered in PDX288 but in this case, it was co-altered with *SOX17.* This co-alteration is distinctive of responder PDXs, with an observed co-occurrence rate of 13.76% in responders and 7.55% in non-responders. *FGFR1* alteration also contributed to the prediction of response. In this case, neither tumor presented known biomarkers of response to MEK inhibition. When treated with Binimetinib, PDX270 was classified as non-responder (PD), as the tumor volume had increased by 144%, even more than in untreated animals (117%). On the contrary, and validating the TCT4U models, PDX288 responded well to treatment (SD), and tumors did not show any significant growth. Interestingly, an integrative genomics screen performed in 229 primary invasive breast carcinomas identified the co-amplification of MYC and the 8p11-12 cytogenetic bands, together with aberrant methylation and expression of several genes spanning the 8q12.1-q24.22 genomic region [58]. This observation coincides with our DCO network derived from whole exome sequencing data, where we could detect the co-amplification of a large cluster of genes located in the 8p11-p12 (*HOOK3*, *FGFR1*) and 8q11.23-q24.22 genomic regions (*TCEA1*, *SOX17*, *CHCHD7*, *NCOA2*, *COX6C*, *MYC*, *NDRG1*) in responder PDXs, but not in non-responder ones (Additional File 1: Table S2, Additional File 3: Figure S3).

Finally, we explored the TCT4U prediction capacity in cytotoxic chemotherapy, where specific oncogenic characteristics should be less related to treatment efficacy. We selected PDX222 and PDX39 to be treated with Paclitaxel. While PDX222 did not present any known biomarker of response, PDX39 sowed an *MCL1* amplification, which has been reported to promote resistance to anti-tubulin chemotherapeutics [14, 59]. Although PDX222 showed alterations that are slightly more common in non-responder than in responder PDXs (*SOX17* and *TP53*, SHAP values −0.177 and −0.208), it also presented an ERBB2 amplification that in our model contributed positively to the final prediction (SHAP value 2.885). Regarding PDX39, *TP53* and *H3F3A* alterations were the main negative contributors to the prediction of response. When treated with paclitaxel, both PDXs showed a progressive disease (PD).

Overall, TCT4U models correctly predicted the outcome of 12 of the 14 treatments tested whereas known biomarkers only predicted correctly 2 of the 14 treatment outcomes. In particular, one of the TCT4U misclassified responses was correctly predicted by known biomarkers, while the rest were either incorrect (4 of 6) or missing (8). In PDXs treated with alpelisib, we correctly classified 3 out of 4 responders and 1 out of 1 non-responders, which makes a balanced accuracy of 0.875. We could not estimate the average balanced accuracy across treatments due to the limited sample size and the imbalanced number of responder and non-responder PDXs per treatment arm. Therefore, the predictions presented and the associated genomic features remain hypothesis in need for further validation.

### Bringing TCT4U from the workbench to the clinics

To explore the clinical potential of TCT4U methodology, we analyzed a cohort of 216 metastatic breast cancer patients being treated at the Memorial Sloan Kettering Cancer Center [41], and for which we have recorded information of their oncogenomic profile and clinical outcome (Additional File 1: Table S6). These metastatic patients had received between 1 and 17 rounds of treatments (median of 2) before being selected for a trial to test a combination of CDK4/6 and aromatase inhibitors. Each tumor was genetically profiled, using the MSK-IMPACT panel, and the clinical outcome of the treatment was recorded as progression free survival (PFS). In this study, one third of the patients did not derive a clinical benefit and relapsed before 5 months. At the other extreme of the distribution, one third of the patients could be treated for more than 10 months and were considered to present a durable clinical benefit. We are aware that a threshold of 10 months might not be relevant in a first line treatment setting, where this drug combination has shown to achieve a median PFS of 24 months [60]. However, the PFS decreases in subsequent lines of therapy and, in a metastatic setting where over half of patients have received prior therapies, a PFS of more than 10 months might still be a good surrogate measure of the clinical benefit.

We did not have PDXs treated with a combination of CDK4/6 and aromatase inhibitors, and the closest TCT4U model for it was derived in response to CDK4/6 inhibition (ribociclib), based on 71 responder and 100 non-responder PDXs. Using this model, only 22.7% patients (49 out of 216) were predicted to respond to treatment and the remaining 77.3% were predicted as non-responders. The majority of patients (78%) relapsed within the first year of treatment but, unfortunately, we have no data in this clinical series as to whether these tumors regressed, at least initially. It thus seems that the outcome measure used to train the TCT4U model (mRECIST), based on relative tumor growth, is not appropriate in most clinical settings.

Without a model for this specific drug combination, and with the aforementioned differences in outcome measures, we decided to adapt our methodology to classify patients based on the duration of the treatment before cancer relapsed. For this, we divided the cohort in three groups and considered the 40 patients for which the tumors relapsed before 4.2 months after the start of the treatment as non-responders, and the 40 for which the time to progression was longer than 9.7 months as responders. The resulting DCO networks for this treatment, which are relatively small compared to TCT4U DCO networks, contain a total of 7 drivers and 6 co-occurrences (see Figure 6A).

**Figure 6.**
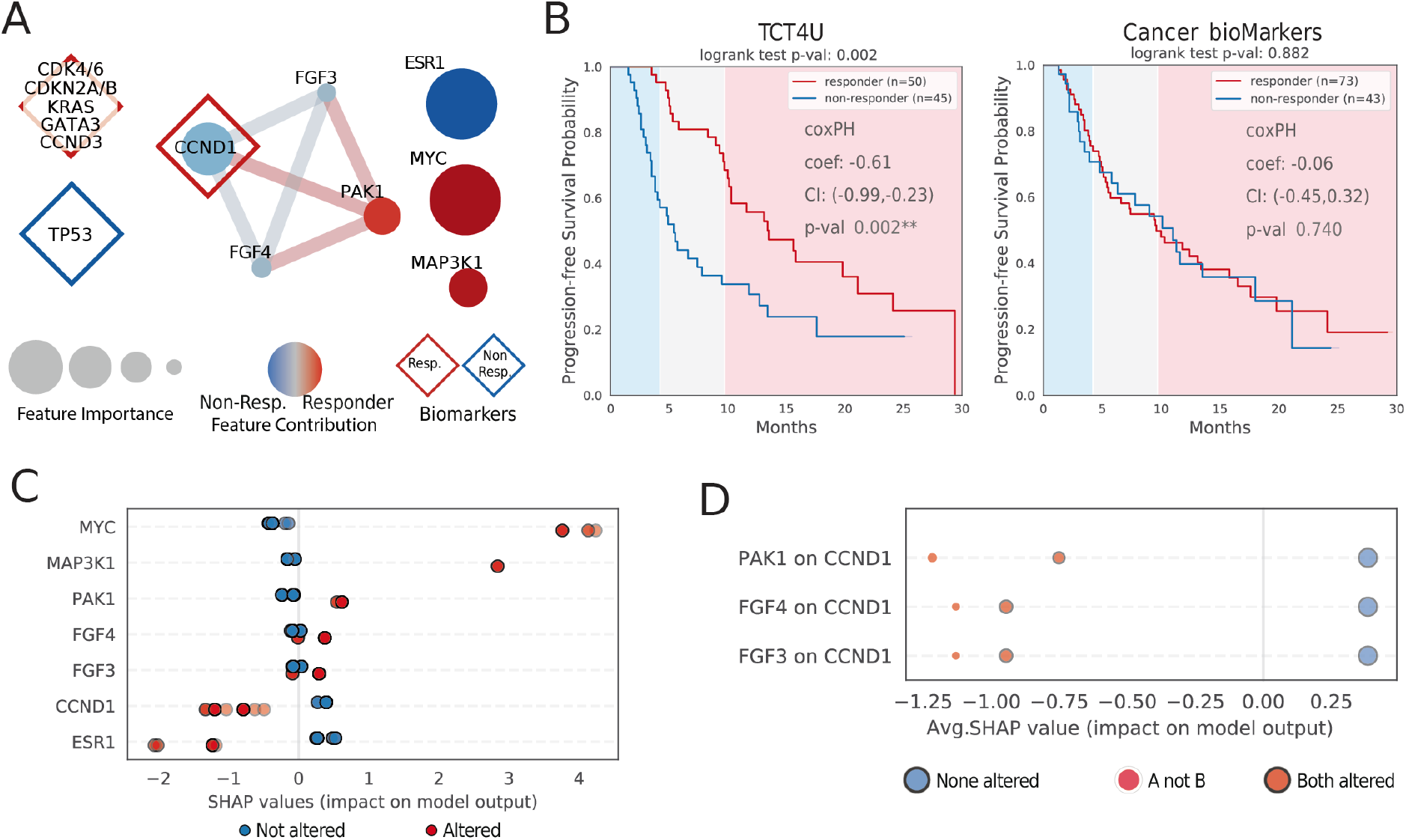
Application of TCT4U to predict treatment outcome in a clinical cohort of HR+/HER2-metastatic breast cancer patients. **(A)** Driver Co-Occurrence networks (DCO) representing the oncogenic alterations and pairs of alterations that are overrepresented in patients that relapsed early (non-responders) or in in patients that derived a durable clinical benefit (responders) from CDK4/6 inhibition combined with an aromatase inhibitor. The size of the nodes represents the average feature importance in the LOOCV. The color of the nodes represents the probability that alterations in a given gene are overrepresented in responders (red) or non-responders (blue). Previously known biomarkers detected in the cohort are annotated with diamond shapes. **(B)** Kaplan-Meier analysis of progression free survival (PFS). TCT4U high-confidence predictions are better able to discriminate between patients that would experience early and late relapse than known biomarkers, with a median time to progression of 5.4 and 13.5 months respectively. **(C)** Summary plot of the SHAP values attributed to individual genes. Each point represents the contribution of a given feature to the prediction of response in a given patient. The color of the points indicates whether a given driver gene was altered or not in each patient. **(D)** SHAP interaction plot showing the positive effect of the co-alteration of *FGF3*, *FGF4*, and *PAK1* with *CCND1* (all in chr11q13-14). The size of the dots is proportional to the number of patients in each category.

We then used the DCO networks to derive the corresponding TCT4U models, which should be able to predict whether a given patient will obtain a significant clinical benefit. In a leave-one-out cross validation, TCT4U models yielded confident scores for 95 out of the 216 patients in the cohort (see *Materials and Methods*). Of these, we predicted that 50 patients would respond and 45 would not respond to the treatment. A Kaplan-Meier analysis of the cross-validation showed that patients predicted to relapse early, with a median time to progression of 5.4 months, derived little clinical benefit compared to the 50 patients predicted to relapse later, whose median time to progression was significantly longer (13.5 months, log-rank test p-value 0.002, Figure 6B). We obtained consistent results when fitting a Cox proportional hazards regression model (correlation coefficient −0.61, p-value 0.002), indicating that TCT4U scores are negatively associated with risk of relapse. The performance of TCT4U models clearly surpasses that of known biomarkers for this drug combination. Although 54% (116 of 216) of patients had at least one annotated biomarker, which is a good coverage compared to other treatments, we could not find a significant association between observed and predicted outcomes, at least in terms of PFS (Figure 6B).

As for the interpretation of the DCO-based model*, MYC* and *MAP3K1* alteration were the strongest explainers of TCT4U predictions of response, with average SAHP values of 3.94 and 2.84 in altered patients. Noteworthy, *MYC* alteration was in common between this model and the ribociclib model derived from PDXs. On the other hand, *ESR1* and *CCND1* were the strongest negative contributors to the predictions, with average SHAP values −1.46 and −0.98; Figure 6C). Oncogenic mutations in *ESR1* are common in metastatic and pretreated breast cancer, emerging as a mechanism of acquired resistance to endocrine therapies that can ultimately result in a lack of response to the combinational therapy [61].

Interestingly, the concomitant alteration of *FGF3* and *FGF4* with *CCND1* (11q13 amplicon) has a positive impact on the SHAP value of *CCND1* altered patients, which goes from −1.13 to −0.94 (Figure 6D). This positive interaction is even stronger when the amplification spans *PAK1* gene (in 11q14), reaching an average SHAP value of −0.75. Indeed, the *FGF3*-*FGF4*-*CCND1* triplet tends to be significantly co-altered with *PAK1* only in patients that relapsed late (p-values of 0.037, 0.038 and 0.027, Additional File 1: Table S2). It is noteworthy that *CCND1* amplification is one of the biomarkers that has been previously associated to response to CDK4/6 blockade. Based on our observations, the genomic context of *CCND1* alteration seems to be relevant in relation to CDK4/6 inhibition and deserves further investigation.

Our results suggest that the proposed methodology could be used to derive DCO networks and train predictive models from the kind of data obtained from interim analyses in oncological clinical trials. Moreover, whenever the time to detect a clinical benefit is reasonable, such the 10 months in this study, TCT4U models could be derived with the first patients and used in population enrichment strategies to establish the bases for new recruitments in adaptive trials.

## Discussion

Cancer sequencing projects have unveiled hundreds of gene alterations driving tumorigenesis, enabling precision oncology. Indeed, current efforts now focus on the analyses of oncogenomic patterns to identify actionable alterations, drugs to modulate them and biomarkers to monitor response. Of particular interest are computational platforms such as OncoKB [13] or the Cancer Genome Interpreter [14], which not only identify oncogenic alterations and potential targets, but also estimate their potential clinical applicability. Most current strategies focus on the identification of a single vulnerability (i.e. driver gene) whose activity can be modulated by a drug. However, given the complexity and heterogeneity in tumors, and the high connectivity between cellular processes, every cancer might respond differently to a certain treatment, depending on its global oncogenomic profile.

Indeed, the analysis of the mutational landscape of cancer has also uncovered the existence of mutual exclusivity and co-occurrence patterns among driver gene alterations [16, 62]. Many computational tools have been developed to identify those combinatorial patterns experimentally (i.e via CRISPR-Cas9 screens [63, 64]) or computationally [65–72]. Patterns of mutual exclusivity can arise from functional redundancy, context-specific dependencies (i.e tumor type or sub-type specific driving alterations), or synthetic lethality interactions. While functional redundancy has been used to reveal unknown functional interactions [72], the synthetic lethality concept has been very successfully applied to the identification of novel therapeutic targets [63, 64] or rational drug combinations [64], and to the prediction of drug response in cell lines [64] and patients [71].

Although less studied, driver co-occurrences are often interpreted as a sign of synergy and in some cases they have shown to be functionally relevant [16–22]. However, they have not yet been exploited for drug response prediction. With the methodology presented in this manuscript, we compared the mutational profiles of tumors that responded or did not respond to a certain drug to define Driver Co-Occurrence (DCO) networks, which capture both genomic structure and putative oncogenic synergy. We then used the DCO networks to train classifiers to identify the best possible treatment for each tumor based on its oncogenomic profile.

The development of tools for personalized treatment prioritization based on genomic profiles is an active field of research. Recently Al-Shahrour and colleagues presented PanDrugs [73], an *in silico* drug prescription tool that uses genomic information, pathway context and pharmacological evidence to prioritize the drug therapies that are most suitable for individual tumor profiles. PanDrugs goes beyond the single-gene biomarker by taking into account the collective gene impact and pathway context of the oncogenic alterations identified in a given patient. However, it combines clinical evidence with *in vitro* drug screening data gathered from cancer cell line panels, which have limited clinical translatability [15, 27, 30, 74].

PANOPLY [75] is another computational framework that uses machine learning and knowledge-driven network analysis approaches to predict patient-specific drug response from multiomics profiles. This tool shows a great potential but the method strongly depends on whole genome and transcriptome patient data, which is not routinely acquired in clinical practice. Other methods like iCAGES [76] have been developed mainly to identify patient-specific driver genes from somatic mutation profiles, which are later used to prioritize drug treatments. However, iCAGES only considers drugs that directly target the identified driver alterations based on current FDA prescription guidelines. All those methods rely on prior knowledge, which is incomplete and biased, and have not been conceived to identify novel co-occurrence patterns from the data and to exploit them for drug response prediction.

With the current implementation of TCT4U, we present a collection of drug-response predictive models for 53 treatments belonging to 20 drug classes, including targeted and more conventional chemotherapies. In a cross-validation setting, our drug-response models attained a global accuracy similar to that of approved biomarkers, but could be applied to twice as many samples, including drug classes for which no biomarker is currently available. Moreover, in an *in vivo* prospective validation, our models correctly predicted 12 out of 14 responses to 5 drugs tested on 13 tumors. However, we could not estimate the average balanced accuracy across treatments due to the limited number of PDXs per treatment arm and the actual imbalance between responder and non-responders.

Obviously, our approach also suffers from some limitations. Due to the lack of systematic reporting of treatment history of the patients enrolled in genomic studies [23], it is difficult to match response to a drug with individual molecular profiles from clinical data. This practically impairs the systematic assessment of the prediction accuracy in patients for computational frameworks like TCT4U, PanDrugs [73], PANOPLY [75], iCAGES [76], or other *in silico* drug prescription tools such as the Cancer Genome Interpreter [14] or OncoKB [13]. Experimental validation of computational approaches is time-intensive and very expensive. Therefore, beyond the thorough experimental validation presented in this manuscript, only PanDrugs and PANOPLY predictions were experimentally validated, although on a single case study performed on a PDX model that was treated with 5 drugs (PanDrugs) or 2 drugs (PANOPLY).

Given the limited clinical representativity of drug screens performed on cell lines [15, 27, 74], we relied on patient-derived xenografts (PDXs) to implement our strategy and to identify biomarkers of drug response. Although PDXs have shown a good level of agreement with the course of disease evolution and treatment response observed in the tumors in the patient [30–33, 77, 78], they present some important drawbacks, such as the eventual loss of intratumoral heterogeneity [79, 80] or certain engraftment bias [30, 81]. Additionally, we have to consider that PDXs might not completely recapitulate the influence of the tissue of origin in tumors that have been implanted subcutaneously in immunodeficient mice and whose stroma has possibly regressed and/or been replaced by mouse stroma, altering thus their sub-clonal evolution and response to treatments [77, 82]. However, our strategy can be readily adapted to derive drug-response models from continuous clinical outcome measures, such as progression free survival, which better represent the data acquired during routine clinical practice and in clinical trials. Indeed, we derived response models on a clinical cohort of breast cancer metastatic patients being treated with a combination of CDK4/6 and aromatase inhibitors, showing a good correlation with progression free survival.

Most importantly, TCT4U drug-response DCO networks are interpretable, and provide clear hints to identify the potential mechanisms of response or non-response present in each tumor. However, one key challenge in interpreting driver alteration co-occurrence patterns is that they can also emerge without necessarily being synergistic if a pair of genes is affected by a common mutagenic process. This commonly happens when several oncogenes are co-amplified as part of the same genomic region and for this reason we have annotated the candidate co-occurring drivers with the genomic location of its constituent genes and distance between them. Moreover, co-occurrence patterns can also emerge as a result of the exposure to other mutagenic processes that increase the mutational burden, the chromosomal instability, or that leave specific mutational signatures [68, 69, 83]. Context or tumor type specific dependencies can also be a source of indirect associations with drug response. Although those confounding factors can obscure the biological interpretation of the DCO networks, they certainly provide valuable information for drug response prediction. Therefore, DCO networks are a valuable asset for hypothesis generation that need to be complemented with orthogonal sources of evidence, and functional validation will always be needed to demonstrate synergy.

We also showed that our methodology is well suited to work with any custom gene panel, provided that the selected genes contribute to the differences in response to the drug being analyzed. As the cost of clinical molecular profiling continues to drop it is very likely that more types of data can be integratively analyzed to improve drug response prediction. However, in order to ensure the clinical translatability of our method in the short term, we decided to focus on well-supported oncogenic alterations that are readily detectable by cost effective methods in the clinical setting. We acknowledge that this is a very conservative decision and we accept that we might be missing biologically relevant information (i.e. non-coding alterations, methylation events or expression changes). Indeed, current clinical biomarkers for patient stratification are mostly based on the detection of histopathological, cytogenetic and immunohistochemical changes that are not always detectable at DNA sequence level. For example, breast cancer patient stratification strategies based on *ER*/*PR* and *ERBB2* status have proven to been very informative, both in terms of prognosis and response to treatment [84]. Accordingly, TCT4U predictions should be regarded as a complementary source of information for clinical decision-making.

## Conclusions

We believe that the computational framework presented, which goes beyond the single gene approach by exploiting co-occurrence patterns, could represent a significant advance towards the development of effective methods for personalized cancer treatment prioritization, with potential applications in population enrichment strategies in the context of adaptive clinical trials. Overall, our strategy represents an opportunity to accelerate the identification and validation of complex biomarkers with the potential to increase the impact of genomic profiling in precision oncology.

## Declarations

### Ethics approval and consent to participate

Mice experiments were conducted following the European Union’s animal care directive (86/609/CEE) and were approved by the Ethical Committee of Animal Experimentation of the Vall d’Hebron Research Institute. Human tumor samples for PDX were obtained after approval from the Ethics Committee of the Vall d’Hebron University Hospital. Written informed consent was signed by all patients. The analysis of patient data was approved by the Memorial Sloan Kettering Cancer Center Institutional Review Board (IRB) and all patients provided written informed consent for tumor sequencing and review of patient medical records for detailed demographic, pathologic, and treatment information. The research conformed to the principles of the Helsinki Declaration.

### Consent for publication

Not applicable.

### Availability of data and materials

Data from the Novartis Institutes for BioMedical Research PDX encyclopedia (NIBR PDXE) is publicly available in the original publication [28] (DOI: 10.1038/nm.3954). Patients’ data is publicly available through the cBioPortal for Cancer Genomics (http://www.cbioportal.org/study?id=breast_msk_2018) and is also deposited in the European Variation Archive (EVA). The accession number for the sequencing data is PRJEB29597, https://www.ebi.ac.uk/eva/?eva-study=PRJEB29597. The accession number for the deposited RNA sequencing data reported in this paper [41] is GEO: GSE122088, https://www.ncbi.nlm.nih.gov/geo/query/acc.cgi?acc=GSE122088. The remaining datasets supporting the conclusions of this article are included within the article (and its additional files).

### Authors’ contributions

L.M., M.D.-F., and P.A. conceived and designed the study. L.M. and M.D.-F developed the methodology and applied it to analyze and interpret the data. A.G-O, M.P., J.A., M.B., and V.S. provided mice model data and helped during its analysis and interpretation. M.S., P.R., and S.C. provided patients’ data and helped during its analysis and interpretation. L.M., M.D.-F., and P.A. wrote the manuscript. V.S., P.R., and S.C. reviewed the manuscript. P.A. supervised the whole study. All authors read and approved the final manuscript.

## Acknowledgements

The authors would like to thank IRB Barcelona and VHIO colleagues for providing feedback on the work, and Dr David Torrents (ICREA-BSC) for critically reading the manuscript. V.S. thanks Faye Su (Novartis Oncology) for providing study reagents alpelisib (BYL719) and ribociclib (LEE011).

## Additional Files

The datasets supporting the conclusions of this article are included in the following additional files:

*Additional File 1 (SupplementaryTables.zip).* This compressed folder contains all the supplementary tables referenced throughout the manuscript, together with their description.

*Additional File 2 (DCO_networks_cytoscape.zip).* This compressed folder contains a cytoscape session for each treatment with all the driver co-occurrence networks generated from whole exome sequencing (WES), IMPACT410 (IM) and Foundation Medicine (FM) gene panels. This information complements the information provided as Additional File 1: Table S2. This file is available at https://doi.org/10.6084/m9.figshare.12789068.v1

*Additional File 3 (SupplementaryFigures.pdf).* This pdf file contains the Supplementary Figures that are referenced throughout the manuscript, together with their figure legends.

Higher resolution figures and extended data are available upon reasonable request.

